# EcoHIV Infection Modulates the Effects of Cocaine Exposure Pattern and Abstinence on Cocaine Seeking and Neuroimmune Protein Expression in Male Mice

**DOI:** 10.1101/2024.04.15.589615

**Authors:** Mark D. Namba, Qiaowei Xie, Kyewon Park, Joshua G. Jackson, Jacqueline M. Barker

**Author notes:** Corresponding Author: Jacqueline M. Barker, Ph.D Department of Pharmacology & Physiology College of Medicine Drexel University 245 North 15th Street, NCB Philadelphia, PA 19102.

## Abstract

Cocaine use disorders (CUDs) and human immunodeficiency virus (HIV) remain persistent public health dilemmas throughout the world. One major hurdle for treating CUD is the increase in cocaine craving and seeking behavior that occurs over a protracted period of abstinence, an effect known as the incubation of craving. Little is known about how HIV may modulate this process. Thus, we sought to examine the impact of chronic HIV infection on the incubation of cocaine craving and associated changes in the central and peripheral immune systems. Here, mice were inoculated with EcoHIV, which is a chimeric HIV-1 construct that produces chronic HIV infection in mice. EcoHIV- and sham-infected mice were conditioned with cocaine daily or intermittently in a conditioned place preference (CPP) paradigm, followed by 1 or 21 days of forced abstinence prior to assessing preference for the cocaine-paired chamber. Under both conditioning regimens, sham mice exhibited incubation of cocaine CPP after 21 days of abstinence. EcoHIV-infected mice conditioned daily with cocaine showed enhanced cocaine seeking at both abstinence timepoints, whereas infected mice conditioned intermittently showed a reversal of the incubation effect, with higher cocaine seeking after 1 day of abstinence compared to 21 days. Analysis of corticolimbic CX3CL1-CX3CR1 and glutamate receptor expression revealed alterations in medial prefrontal cortex (mPFC) CX3CL1 and nucleus accumbens (NAc) GluN2A receptors that correlated with cocaine seeking following daily cocaine exposure. Moreover, examination of peripheral immune markers showed that the effect of abstinence and EcoHIV infection on these measures depended on the cocaine exposure regimen. Altogether, these results highlight the importance of cocaine abstinence and exposure pattern as critical variables that modulate HIV-associated neuroimmune outcomes and relapse vulnerability.

## Introduction

Use of psychostimulants such as cocaine remains a significant public health concern worldwide. This is especially true among people living with human immunodeficiency virus (HIV; PLWH) who use drugs at a higher rate than the general population and may face unique challenges in managing drug use (Avants et al., 1998; Durvasula and Miller, 2014; Krishnan et al., 2018). Broadly, substance use can increase the risk of HIV transmission through risky sexual behaviors, sharing of injection equipment, and difficulties in managing daily antiretroviral therapy (ART) (Baum et al., 2009; Heaton et al., 2010; Lucas, 2011; Kumar et al., 2015; Hartzler et al., 2017; Shiau et al., 2017). Many studies have demonstrated that addictive drugs, such as cocaine, can facilitate the pathophysiology of HIV (Baum et al., 2009; Purohit et al., 2011; Dahal et al., 2015; Dash et al., 2015). However, far fewer studies have attempted to address how chronic HIV infection may impact addiction-related clinical outcomes. This is particularly important given that successful treatment of substance use disorders may greatly improve the clinical management of HIV among people who use drugs (Durvasula and Miller, 2014; Dash et al., 2015; Kumar et al., 2015; Bertholet et al., 2023).

Relapse to cocaine use remains a major challenge for individuals with cocaine use disorder (CUD) seeking long-term abstinence. One key factor that contributes to drug relapse is exposure to drug-associated environmental contexts or cues which can trigger intense cravings and drug-seeking behavior. This is evident – and even potentiated - after prolonged periods of abstinence from drug use (Childress et al., 1999; Parvaz et al., 2016). Many preclinical studies have modeled the potentiation of drug seeking in rodent models of cocaine-seeking behavior following protracted abstinence, suggesting that this ‘incubation’-like effect occurs across species. This has enabled identification of corticolimbic neural substrates that, at least in part, mediate this process (Tran-Nguyen et al., 1998; Grimm et al., 2001; Conrad et al., 2008; Pickens et al., 2011; Li et al., 2015; Lubbers et al., 2016). Such substrates include AMPA and NMDA glutamate receptors within corticostriatal circuits, which a plethora of studies have implicated in abstinence-induced facilitation of cocaine relapse-like behavior (Conrad et al., 2008; Van Den Oever et al., 2008; Kalivas, 2009; Li et al., 2015; Koob and Volkow, 2016). In addition to drug-induced glutamatergic plasticity, addictive drugs and HIV can impair neuromodulatory immune function within corticolimbic reward circuitry that can contribute to drug-seeking behavior (Namba et al., 2021). Fractalkine (i.e., CX3CL1-CX3CR1) signaling is a critical neuroimmune mechanism that regulates crosstalk between neurons and microglia, the brain’s resident immune cells and the principal cellular reservoir for HIV in the brain. While some studies indicate that changes in corticolimbic fractalkine signaling are associated with cocaine seeking behavior (Montesinos et al., 2020; Rosa et al., 2022), other studies suggest a possible protective role of fractalkine signaling in the context of HIV (Suzuki et al., 2011; Duan et al., 2014). Importantly, fractalkine signaling can negatively regulate glutamatergic excitation through inhibition of AMPA receptor-mediated EPSCs and by enhancing astrocytic glutamate transport via GLT-1 (Paolicelli et al., 2014; Lauro, 2015). Taken together, these studies suggest that corticostriatal glutamate-neuroimmune interactions may underlie HIV-induced changes in drug-seeking behavior. However, no studies to date have identified drug-induced glutamatergic and neuroimmune adaptations within corticolimbic circuitry that are modulated by chronic HIV infection and other important and clinically relevant factors such as drug exposure pattern. Thus, the present study sought to dissect these crucial variables in a model of incubated cocaine craving, in which abstinence-dependent corticostriatal glutamatergic and neuroimmune adaptations are known to occur (Conrad et al., 2008; Li et al., 2015; Kim et al., 2018; Namba et al., 2021).

Across rodent models, HIV and its protein products can alter drug-seeking behavior and associated neuronal plasticity (Namba et al., 2023b). For example, the HIV transgenic rat, which systemically and constitutively expresses 7 of the 9 HIV-1 genes (Reid et al., 2001), exhibits corticolimbic synaptic dysfunction, neuroinflammation, altered reward seeking behavior, and other learning impairments (Vigorito et al., 2007; Royal et al., 2012; Moran et al., 2014; McIntosh et al., 2015; Reid et al., 2016; Wayman et al., 2016; McLaurin et al., 2018; de Guglielmo et al., 2020). Other models, such as transgenic mice that constitutively or inducibly express HIV-1 proteins such as trans-activator of transcription (Tat) or the envelope protein gp120, exhibit similar reward learning impairments as well as synaptic and neuroimmune dysfunction (Kesby et al., 2014, 2017; Paris et al., 2014a, 2014b; Hoefer et al., 2015; Dickens et al., 2017; Schier et al., 2017; Thaney et al., 2018). While these models have provided invaluable insights into the interaction between HIV infection and drug use, they do not fully recapitulate the complexity of chronic HIV infection in humans, which constrains the translational potential of preclinical studies employing such models. The EcoHIV mouse model, which uses a chimeric virus to infect mice with a modified HIV-1 genome, has been developed to address some of these limitations by permitting the examination of how chronic viral infection, as opposed to viral protein exposure alone, impacts neurocognitive function. In this model, the gp120 envelope protein is replaced with gp80 from the ecotropic murine leukemia virus, which permits successful HIV-1 infection and subsequent expression of HIV-1 genes in mice (Potash et al., 2005). EcoHIV-infected mice exhibit transient viremia during the first week of infection and maintain immunocompetency amidst low levels of chronic infection for months following initial exposure (Gu et al., 2018). Concurrently, these mice also exhibit spatial learning impairments that are not rescued by ART treatment following the establishment of chronic EcoHIV infection (Gu et al., 2018; Kelschenbach et al., 2019). Given that PLWH can experience cognitive impairments despite undetectable viral load (Heaton et al., 2010, 2011), these preclinical findings highlight the translational value of this model and its utility in studying the neurobehavioral sequelae of comorbid HIV and CUDs. Nevertheless, findings from clinical studies examining the interactive effects of HIV and drug use on cognitive function and behavior are mixed, with numerous discrepancies across studies that may be due to factors such as drug use patterns, polysubstance use, ART type, socioeconomic factors, among many others (Norman et al., 2009). This further highlights the need for experimentally controlled preclinical examination of HIV and drug interactions on cognition and behavior. Thus, we employed the EcoHIV model to probe the impact of varying durations of cocaine abstinence and of exposure pattern on HIV-induced deficits in cocaine seeking and associated neuroimmune adaptations.

In this manuscript, we investigate the neurobehavioral effects of prolonged abstinence from cocaine on conditioned place preference (CPP) within the EcoHIV mouse model. We show that prolonged abstinence from cocaine, regardless of exposure pattern, increases cocaine seeking behavior, while the potentiating effect of EcoHIV infection on this process depends on the exposure pattern. We also demonstrate that cocaine abstinence- and EcoHIV-induced peripheral immune alterations also depend on the pattern of cocaine exposure. Within the mPFC and NAc, which are key corticolimbic brain regions that are sensitive to HIV infection and mediate cocaine-seeking behavior, we measured expression levels of fractalkine and its receptor, CX3CR1, in addition to AMPA and NMDA glutamate receptor subunits. Here, we demonstrate that the impact of EcoHIV infection on fractalkine signaling system and glutamate receptor expression, and its relationship to cocaine seeking, is brain region- and exposure pattern-specific. Our findings collectively suggest that the frequency of cocaine exposure is an important factor that modulates relapse susceptibility and associated neurobiological adaptations after prolonged abstinence within the context of HIV, providing insight into the development of effective relapse prevention strategies for PLWH.

## Methods

### Animal subjects

108 male C57BL/6J mice were obtained from The Jackson Laboratory (Bar Harbor, ME, USA) at 9 weeks old. Mice were co-housed in groups of 2-4 and remained in their home cages for 7 weeks from date of arrival following EcoHIV inoculation (see below) and prior to behavioral testing within a temperature- and humidity-controlled vivarium on a 12:12 hr light:dark cycle. Once per week on weeks 1, 3, and 5 post-inoculation, mice were briefly anesthetized with 5% isoflurane to collect submandibular blood for use in future studies. Mice had *ad libitum* access to food and water for the entire duration of the study. All experiments were approved and performed in accordance with the Institutional Animal Care and Use Committee of Drexel University and the National Institutes of Health’s *Guide for the Care and Use of Laboratory Animals*. **Figure 1** depicts a timeline of experimental procedures.

**Figure 1.**
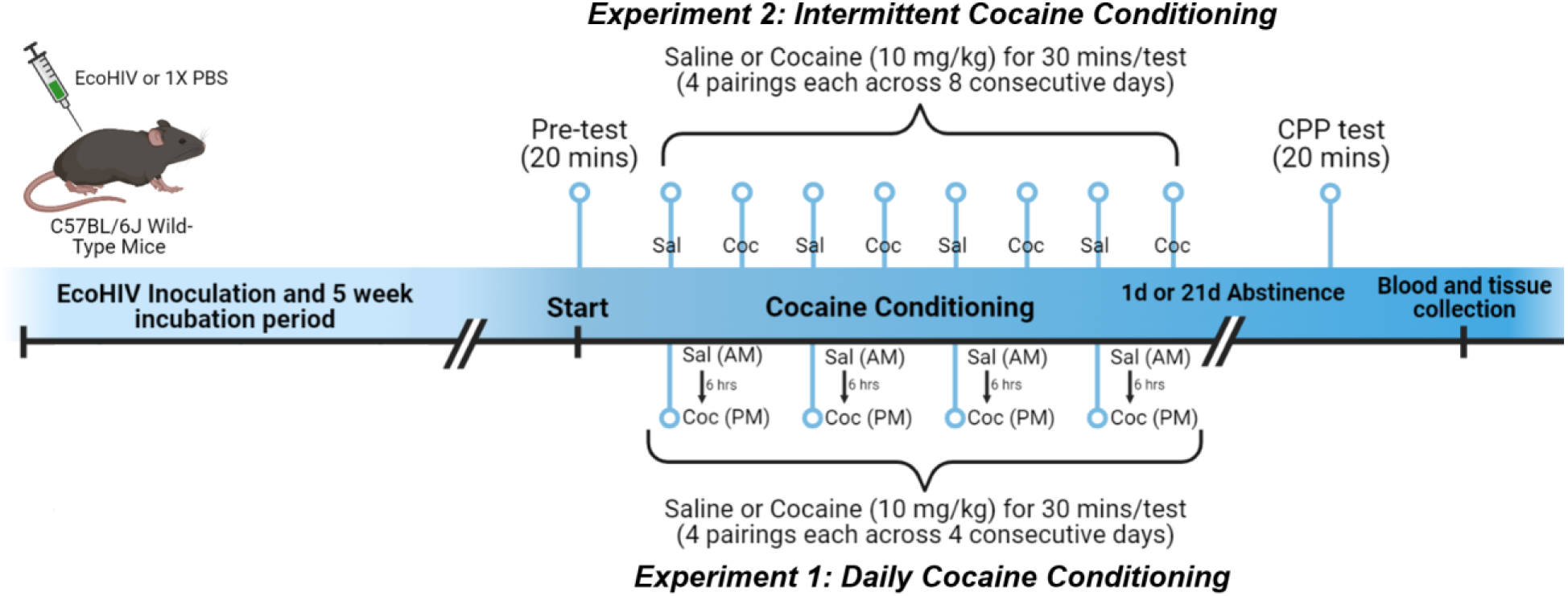
Timeline of experimental procedures. Male C57BL/6J mice (N=108) were inoculated with EcoHIV (300-400 ng p24, i.p.) or 1X PBS, followed by blood collection on weeks 1, 3, and 5 post-inoculation. Beginning on Week 6 post-inoculation, mice were tested for their CPP chamber bias during a 20-min pretest session. Using a biased CPP design during cocaine conditioning, cocaine injections (10 mg/kg, i.p.) were paired with the chamber that was least preferred during the pretest while saline injections (1 mL/kg, i.p.) was paired with the opposite chamber. **In Experiment 1**, mice received a saline pairing in the morning, followed by a cocaine pairing 6 hours later, for a total of 4 pairings each as in Experiment 1 (i.e., “daily conditioning”). **In Experiment 2**, cocaine and saline pairings (4 each) occurred on alternating days (i.e., “intermittent conditioning”). After cocaine conditioning, mice were placed in forced abstinence for 1 or 21 days prior to a CPP test, where chamber preference was assessed in a drug-free state. Mice were sacrificed immediately after CPP testing for tissue collection.

### Drugs & reagents

Cocaine hydrochloride (NIDA Drug Supply Program, RTI International, Research Triangle Park, NC, USA) was dissolved in sterile saline to a concentration of 1 mg/mL immediately prior to cocaine conditioning and was administered at a dose of 10 mg/kg, i.p. On the day of EcoHIV inoculations, virus was diluted in sterile 1X phosphate-buffered saline (1X PBS) and administered to mice at a final concentration of 300-400 ng of HIV- 1 p24 per 0.2 mL PBS.

### EcoHIV synthesis, inoculation, and validation of infection status

Plasmids for EcoHIV-NDK (kindly provided by Dr. David Volsky, Mount Sinai Icahn School of Medicine) were purified from bacterial stocks (Stbl2 cells, ThermoFisher #10268019) using an endotoxin-free purification kit (ZymoPure #D4200). DNA was transfected into nearly confluent (80-90%) 10 cm^2^ plates of low-passage LentiX 293T (Takara #632180) using a calcium phosphate transfection protocol, and supernatants were collected at 48 hrs post-transfection. Cellular debris were pelleted by centrifugation (1500 x g) on a benchtop centrifuge at 4°C followed by passage through a 40 µm cell strainer. Supernatant containing viral particles was mixed 4:1 with a lentiviral concentrator solution (4x; MD Anderson) comprised of 40% (w/v) PEG-8000 and 1.2M NaCl in PBS (pH 7.4). This preparation incubated overnight at 4°C on an orbital shaker (60 rpm) then centrifuged at 1500 x g for 30 mins at 4°C. The resulting pellet was rinsed 1x with PBS then centrifuged again at 1500 x g for 5 mins. The solution was removed, and the pellet was then resuspended in cold, sterile PBS. Viral titer (p24 core antigen content) was determined initially using a LentiX GoStix Plus titration kit (Takara #631280) and subsequently by HIV p24 AlphaLISA detection kit (PerkinElmer #AL291C). Viral stocks were aliquoted and stored at -80°C until future use.

One week after arriving at the animal facility, mice were briefly anesthetized with 5% isoflurane and inoculated with EcoHIV (0.2 mL; n=48) or 1X PBS (i.e., sham; 0.2 mL; n=48). To validate infection status, DNA was isolated from 10 mg punches of frozen spleens, collected at the end of the study (see below), using the Qiagen QIAamp DNA Mini Kit (Cat #51304) according to manufacturer instructions. DNA samples were evaluated for HIV-1 LTR DNA by the University of Pennsylvania Center for AIDS Research (CFAR). The primer and probe sequences used were as follows:

F: 5’-GCCTCAATAAAGCTTGCCTTGA-3’

R: 5’-GGGCGCCACTGCTAGAGA-3’

FAM/BHQ probe: 5’-CCAGAGTCACACAACAGACGGGCACA-3’

OD values for each sample were obtained using a NanoDrop™ spectrophotometer (Thermo Scientific) and used to estimate the input cell numbers to normalize the final data. HIV-1 DNA was undetectable in all 12 of the representative sham samples tested and 2 mice inoculated with EcoHIV that had undetectable levels of HIV-1 DNA were excluded from the study.

### Cocaine conditioned place preference

Five days after the final submandibular blood collection, mice were placed within a three-chambered CPP apparatus (Med Associated Inc., St. Albans, VT, USA) that consisted of a gray central chamber with smooth flooring, a black chamber with bar flooring, and a white chamber with grid flooring. Here, mice were placed in the center chamber and their initial side bias was assessed during a 20-min pretest session. A biased cocaine conditioning design was implemented such that cocaine was always paired with the least-preferred chamber. For *Experiment 1*, mice received two 30-min conditioning sessions per day, beginning with a saline pairing in the morning followed by a cocaine pairing in the afternoon 6 hours later. This was repeated for 4 consecutive days for a total of 8 pairings (4 saline and 4 cocaine). For *Experiment 2*, mice received one 30- min conditioning session per day, beginning with a saline pairing on day 1, followed by a cocaine pairing on day 2. This intermittent pattern of cocaine exposure continued for a total of 8 consecutive days, resulting in 8 total pairings (4 saline and 4 cocaine). Throughout cocaine conditioning, total locomotor activity was tracked during each conditioning session. After cocaine conditioning, mice entered a period of forced abstinence for either 1 or 21 days, where they remained undisturbed in their home cages. Following forced abstinence, mice were placed back into the CPP apparatus for a 20-min test session in a drug-free state, in which preference for the cocaine-paired chamber was assessed.

### Tissue collection & processing

Immediately after the post-abstinence CPP test, mice were deeply anesthetized with isoflurane and submandibular blood was collected prior to rapid decapitation. Briefly, mice were induced with 5% isoflurane and submandibular blood was collected using an 18-gauge needle and stored over ice in 0.5 mL tubes containing EDTA (Greiner, Monroe, NC, USA). Fresh whole brains were flash frozen in 2-methylbutane and whole spleen tissue was frozen over dry ice. Blood was centrifuged at 8700 x g for 20 mins at 4°C and the plasma supernatant was collected in PCR-clean microcentrifuge tubes and stored at -80°C. Frozen brains were dissected over dry ice and biopsy punches were taken from the mPFC and NAc. Tissue punches were crudely homogenized with a plastic mortar followed by sonication in 100-250 uL of RIPA lysis buffer containing protease and phosphatase inhibitors (Santa Cruz Biotechnology, Dallas, TX, USA). Tissue homogenates were then centrifuged at 12,000 x g for 10 mins and supernatants were collected. Total protein concentrations were determined using the Pierce BCA Protein Assay Kit (Thermo Fisher, Waltham, MA, USA) and stored at -80°C for future analyses. *Western blot analyses*

Total protein from whole cell lysates of mPFC and NAc (6-8 µg) were loaded onto NuPAGE 4-12% Bis-Tris gels (Invitrogen) under reduced and denatured conditions (Pierce 4X LDS sample buffer; 10X NuPAGE Sample Reducing Agent). Protein was transferred to nitrocellulose membranes under semi-dry conditions (Invitrogen iBlot 2) and membranes were blocked with 5% milk in 1X tris-buffered saline + 0.1% Tween-20 (TBST). Membranes were probed overnight at 4°C in primary antibody containing blocking buffer, followed by 6 x 5 min washes in blocking buffer. Membranes were then probed with secondary antibody in 1X TBST for 1 hour and then washed 6 x 5 mins in 1X TBST. Chemiluminescent activation of the membranes was achieved using the SuperSignal™ West Pico PLUS Chemiluminescent Substrate and chemiluminescence was measured using a Licor Odyssey® FC imager. Optical density for each protein band was analyzed using ImageJ. Protein expression for each band of interest was normalized to an internal loading control that was used across all gels, and these adjusted values were normalized to that of GAPDH as an internal control for each gel lane.

### Plasma cytokine, chemokine, and growth factor analyses

Plasma collected at sacrifice was partially diluted with 1X PBS + 0.1% Triton X- 100 and cytokine, chemokine, and growth factor expression levels were determined using the Mouse Cytokine/Chemokine 32-Plex Discovery Assay® Array (MD32; Eve Technologies, Calgary, AB Canada). Each sample was analyzed in duplicate and the average of each pair of readings was used as the final measure for each sample across all targets. Plasma collected from a separate cohort of sham-inoculated mice, exposed to 8 saline pairings akin to mice in the CPP experiments, was used as a reference control group.

### Statistical analyses

Locomotor behavior across cocaine and saline pairings was analyzed via three-way repeated measures ANOVAs, with Pairing (repeated measure), Drug, and Infection status as factors. Cocaine CPP scores were calculated as time spent in the cocaine-paired chamber during the pretest subtracted from this time during the post-abstinence CPP test (i.e., “Post-Pre”). This calculation was conducted across four 5-min bins for the 20-minute pre- and post-test assessments, which were then analyzed via three-way repeated measures ANOVAs, with Bin (repeated measure), Infection Status, and Abstinence as factors. For all behavior analyses, the Holm-Šídák method was used to correct for multiple *post hoc* comparisons. To examine the effect of abstinence and cocaine exposure pattern on HIV-1 viral DNA levels in spleens, a two-way ANOVA was used, with Abstinence and Exposure Pattern as factors, and the Holm-Šídák method was applied to correct for multiple *post hoc* comparisons here. To assess mPFC and NAc protein expression, two-way ANOVAs were used to assess group mean differences for each experiment, with Infection Status and Abstinence as factors. Simple linear regressions were also used to calculate the correlation between cocaine CPP scores and protein expression. The Holm-Šídák method was used to adjust the p-values and correct for multiple testing. Only analyses that survived this p-value correction were considered significant at α = 0.05 significance level. Plasma cytokine expression was analyzed via one-way ANOVAs, with saline-treated sham controls as a reference group. Bartlett’s tests were used to test for unequal variances; in such cases, a Welch’s ANOVA test was applied. Again, the Holm-Šídák method was used to adjust the p-values for multiple testing, and Dunnett’s tests for multiple comparisons were used to further probe individual group differences, relative to the sham-saline control group, for analyses that survived this correction. The ROUT method was employed, with Q coefficient = 0.1%, to screen for outliers. Across repeated measures analyses, Greenhouse-Geiser corrections were applied where necessary to account for lack of sphericity. n = 2 EcoHIV-inoculated mice were excluded from the study due to undetectable HIV DNA. For each analysis, the significance threshold was set at α = 0.05. All analyses were conducted using GraphPad Prism v.10.

## Results

### Experiment 1: Daily cocaine conditioning and place preference

We first tested the impact of EcoHIV infection on the incubation of cocaine seeking behavior in mice conditioned daily with cocaine (**Figures 2A, B**). A repeated measures three-way ANOVA of locomotor behavior (**Figure 2A**) during cocaine conditioning revealed significant main effects of Pairing (*F*_(3,138)_ = 8.42, p < 0.0001), Infection (*F*_(1,46)_ = 9.093, p = 0.0042), and Drug (*F*_(1,46)_ = 160.8, p < 0.0001), as well as significant Pairing X Infection (*F*_(3,138)_ = 7.561, p = 0.0001), Pairing X Drug (*F*_(3,138)_ = 8.897, p < 0.0001), Infection X Drug (*F*_(1,46)_ = 12.9, p = 0.0008), and Pairing X Drug X Infection (*F*_(3,138)_ = 6.0, p = 0.0007) interactions. *Post-hoc* analyses of the three-way interaction revealed a significant increase in locomotor activity among EcoHIV-infected mice relative to shams for pairings 3 and 4.

**Figure 2.**
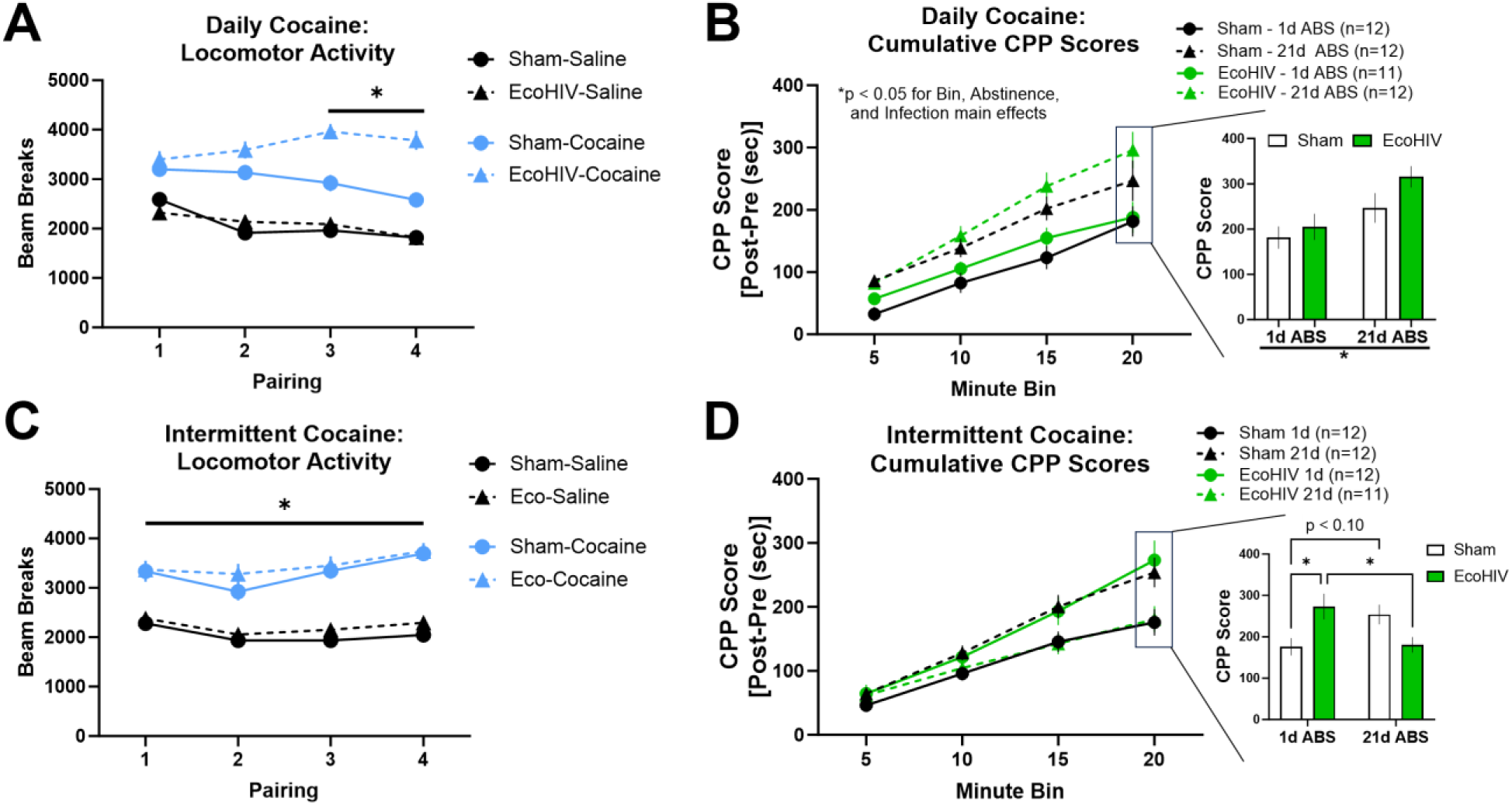
Cocaine-induced locomotion and conditioned place preference. **(A)** In Experiment 1 (daily conditioning), cocaine increased locomotor activity relative to saline treatment across all 4 pairings, and EcoHIV-infected mice exhibited potentiated locomotor behavior on pairings 3 and 4 (*p < 0.05 comparing EcoHIV-Cocaine vs. Sham-Cocaine; n = 23-24/group). **(B)** Among shams, cocaine seeking was increased across the session following 21d ABS relative to 1d ABS. This was further potentiated by EcoHIV infection at each timepoint (*p < 0.05 for main effect of Abstinence). **(C)** In Experiment 2 (intermittent conditioning), cocaine similarly produced a greater locomotor response relative to saline (*p < 0.05 comparing Sham- or EcoHIV-cocaine to Sham- or EcoHIV-saline for each pairing; n = 23-24/group). However, EcoHIV infection did not alter locomotor behavior relative to shams at each pairing. **(D)** Among mice conditioned intermittently, sham mice showed a strong trend towards abstinence-induced potentiation of cocaine seeking, while EcoHIV-infection produced the reverse effect (p < 0.0001 for Bin x Abstinence x Infection interaction from three-way ANOVA; *post hoc* comparisons at each timepoint did not achieve statistical significance). *p < 0.05. Error bars = ±SEM.

Mice were then tested for cocaine CPP after 1 or 21 days of forced abstinence (**Figure 2B**). CPP scores were calculated as follows: [Posttest time spent in cocaine-paired chamber] – [Pretest time spent in cocaine-paired chamber]. This calculation was conducted for each 5-min bin to examine cocaine CPP across the 20-min session. A repeated measures three-way ANOVA revealed significant main effects of Bin (*F*_(1.503,64.63)_ = 158.4, p < 0.0001), Infection (*F*_(1,43)_ = 4.26, p = 0.0451), and Abstinence (*F*_(1,43)_ = 10.19, p = 0.0026). Here, both sham- and EcoHIV-infected mice exhibited incubation of cocaine CPP after 21 days of forced abstinence and EcoHIV infection further increased cocaine CPP at both abstinence timepoints relative to sham animals. A two-way ANOVA of the total CPP scores revealed a significant main effect of Abstinence (*F*_(1,43)_ = 10.16, p = 0.0027), but no significant main effect of Infection or interaction effect. Altogether, these results indicate that daily exposure to cocaine in EcoHIV infected mice potentiated cocaine-induced locomotion and cocaine-seeking behavior across the test session for both abstinence timepoints.

### Experiment 2: Intermittent cocaine conditioning and place preference

We next tested the modulatory effect of EcoHIV infection on the incubation of cocaine seeking in mice conditioned with cocaine in an alternating (or “intermittent”) manner (**Figure 2C,D**). A repeated measures three-way ANOVA of locomotor activity (**Figure 2C**) during cocaine conditioning revealed significant main effects of Pairing (*F*_(3,138)_ = 12.55, p < 0.0001) and Drug (*F*_(1,46)_ = 149.2, p < 0.0001), as well as a significant Pairing X Drug interaction (*F*_(3,138)_ = 6.178, p = 0.0006). A *post hoc* analysis of this interaction revealed a significant cocaine-induced increase in locomotor behavior for each conditioning session, regardless of infection status.

After 1 or 21 days of forced abstinence following cocaine conditioning, mice were tested for cocaine CPP in a drug-free state (**Figure 2D**). A repeated measures three-way ANOVA of cocaine CPP revealed a significant main effect of Bin (*F*_(1.507,63.31)_ = 223.1, p < 0.0001), as well as significant Infection X Abstinence (*F*_(1,42)_ = 7.892, p = 0.0075) and Bin X Infection X Abstinence (*F*_(3,126)_ = 12.36, p < 0.0001) interactions. Despite the significant interaction, *post hoc* analyses revealed no significant group differences at each bin. A two-way ANOVA of the total CPP scores revealed a significant Infection X Abstinence interaction (*F*_(1,43)_ = 12.68, p = 0.0009), and *post hoc* tests revealed a significant increase in cocaine seeking due to EcoHIV infection at 1d ABS (*p < 0.05), a significant reduction in cocaine seeking among EcoHIV-infected mice from 1d ABS to 21d ABS (*p < 0.05), and a trend towards potentiated cocaine seeking from 1d ABS to 21d ABS among sham mice (p < 0.10). Taken together, EcoHIV-infected mice that underwent 1 day of forced abstinence exhibited potentiated cocaine CPP relative to shams. However, EcoHIV-infected mice exhibited reduced cocaine seeking after protracted (21d) abstinence relative to 1d ABS.

### HIV-1 DNA in spleen tissue

To validate the infection status of mice from Experiments 1 and 2, we isolated and purified DNA from the spleens of sham- and EcoHIV-infected mice and quantified HIV-1 long terminal repeat (LTR) DNA levels via PCR (**Figure 3**). Two (out of 48) EcoHIV-inoculated mice were excluded from the study, as they did not exhibit any detectable levels of viral DNA. A two-way ANOVA revealed significant main effects of Abstinence (*F*_(1,42)_ = 10.37, p = 0.0025), Exposure Pattern (*F*_(1,42)_ = 21.13, p < 0.0001), and a significant Abstinence X Exposure Pattern interaction (*F*_(1,42)_ = 8.462, p = 0.0058). *Post hoc* tests revealed that HIV-1 LTR DNA levels were significantly greater in mice conditioned daily after 1d ABS compared to all other groups. Notably, no differences were observed between the other three groups. These data suggest that cocaine exposure pattern critically determined the impact of acute cocaine abstinence on viral DNA burden in spleen.

**Figure 3.**
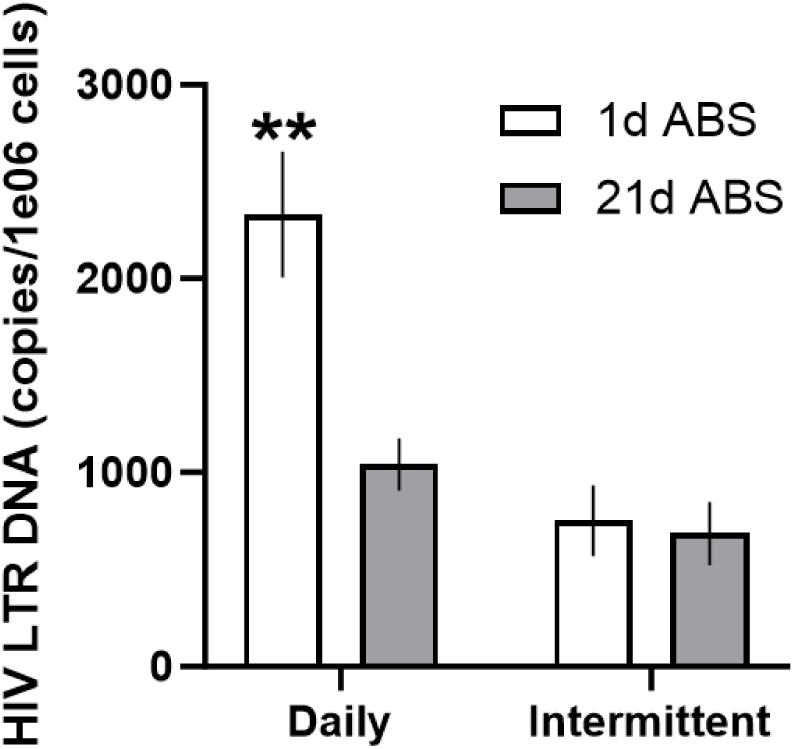
Splenic HIV-1 LTR DNA levels following daily or intermittent cocaine conditioning and 1 or 21 days of abstinence. HIV-1 LTR DNA was evaluated from all EcoHIV-infected mice. Two mice that had undetectable levels of HIV-1 DNA, akin to all sham samples tested, were excluded from the study. Among mice exposed to cocaine daily, splenic HIV-1 LTR DNA levels were increased at 1 day (1d ABS) compared to the 21 days of abstinence (21d ABS). Conversely, no differences were observed in DNA levels between abstinence timepoints among mice exposed to cocaine intermittently. n = 11- 12/group, **p < 0.01 relative to all other groups. Error bars = ±SEM.

### mPFC and NAc glutamate receptor and fractalkine system expression

Given the putative role of the mPFC and its glutamatergic projections to limbic regions such as the NAc in mediating cocaine seeking, as well as the susceptibility of both regions to drug- and HIV-induced neuroimmune dysfunction, we examined whether EcoHIV infection, cocaine exposure pattern, and abstinence uniquely altered mPFC and NAc fractalkine signaling system and glutamate receptor expression. Thus, we examined the expression of fractalkine, CX3CR1, AMPA receptor subunits GluA1 and GluA2, and NMDA receptor subunits GluN2A and GluN2B. For Experiment 1, analysis of mPFC protein expression revealed no significant main effects of Infection or Abstinence, nor a significant interaction, on protein expression following p-value correction. However, mPFC fractalkine expression was positively correlated with cocaine CPP scores among sham- but not EcoHIV-infected mice despite no group mean differences (**Figure 4**; R^2^ = 0.4066, *F*_(1,21)_ = 14.39, p = 0.0011, p-adj. = 0.0131). For Experiment 2, two-way ANOVAs of mPFC protein expression, following p-value correction for multiple testing, revealed no significant effect of abstinence or EcoHIV infection on any of the measured proteins. Similarly, simple linear regression analyses of mPFC protein expression and cocaine CPP, following p-value correction, revealed no significant correlations among sham- or EcoHIV-infected mice for any protein targets.

**Figure 4.**
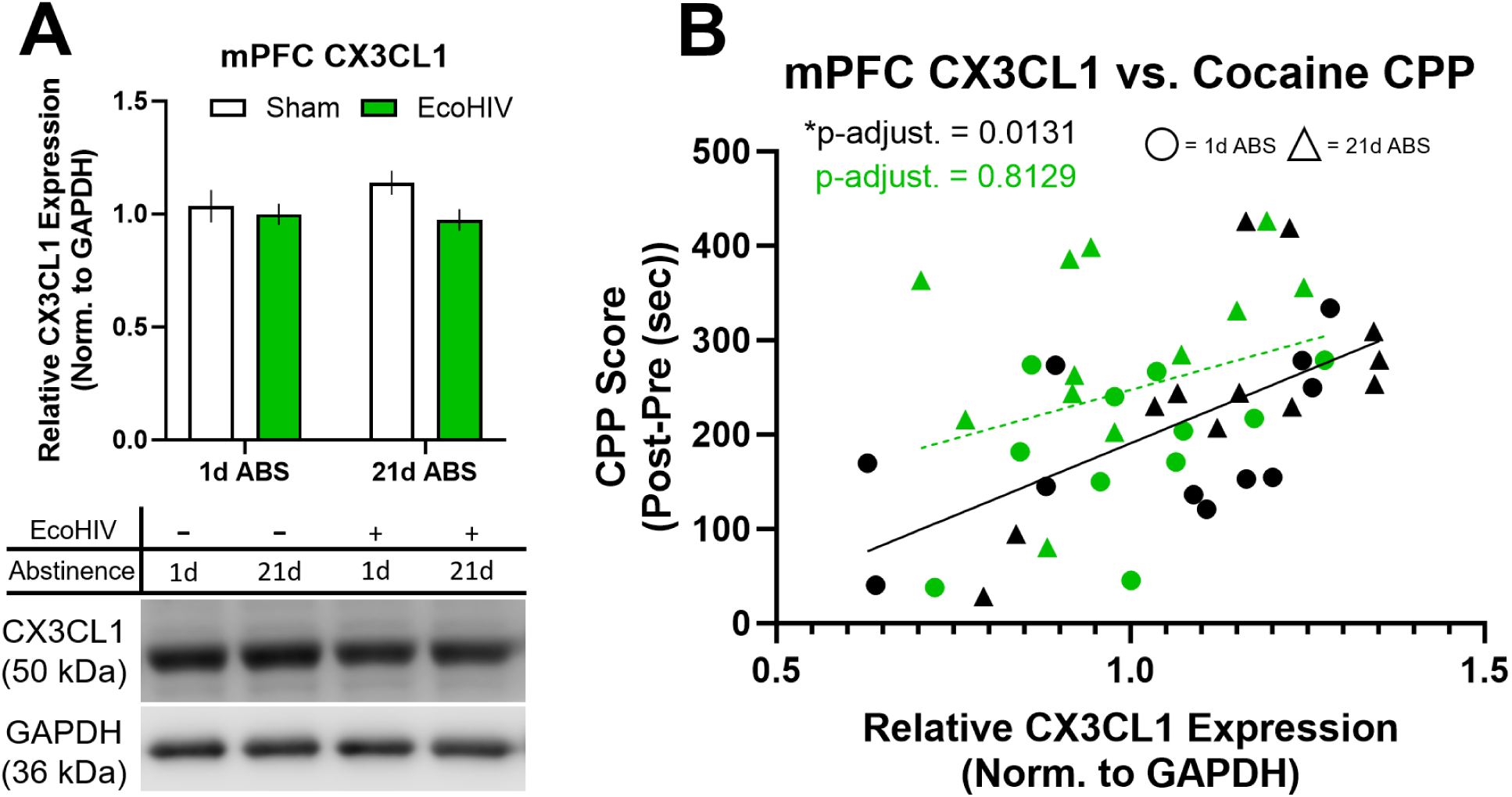
mPFC CX3CL1 expression and associated cocaine seeking following daily conditioning and abstinence. (**A**) Among mice conditioned daily, no group mean differences in mPFC CX3CL1 expression were detected due to EcoHIV, abstinence, or their interaction. Error bars = ±SEM; n = 11-12/group. (**B**) However, cocaine seeking behavior was significantly correlated with CX3CL1 expression only among sham mice across abstinence time points.

Two-way ANOVAs examining mean differences in NAc protein expression did not reveal any significant effect of abstinence, EcoHIV infection, or their interaction across any of the measured targets, for either experiment, following p-value correction. However, in Experiment 1, simple linear regression analyses revealed a significant positive correlation between NAc GluN2A expression and cocaine CPP scores among sham- but not EcoHIV-infected mice (**Figure 5**; R^2^ = 0.3333, *F*_(1,21)_ = 11.84, p = 0.0025, p-adj. = 0.0296). Taken altogether, these results indicate that cocaine seeking following daily, repeated exposure is associated with mPFC fractalkine and NAc GluN2A expression in the absence of EcoHIV infection. This may implicate alternative mechanisms in the regulation of EcoHIV-induced cocaine seeking deficits.

**Figure 5.**
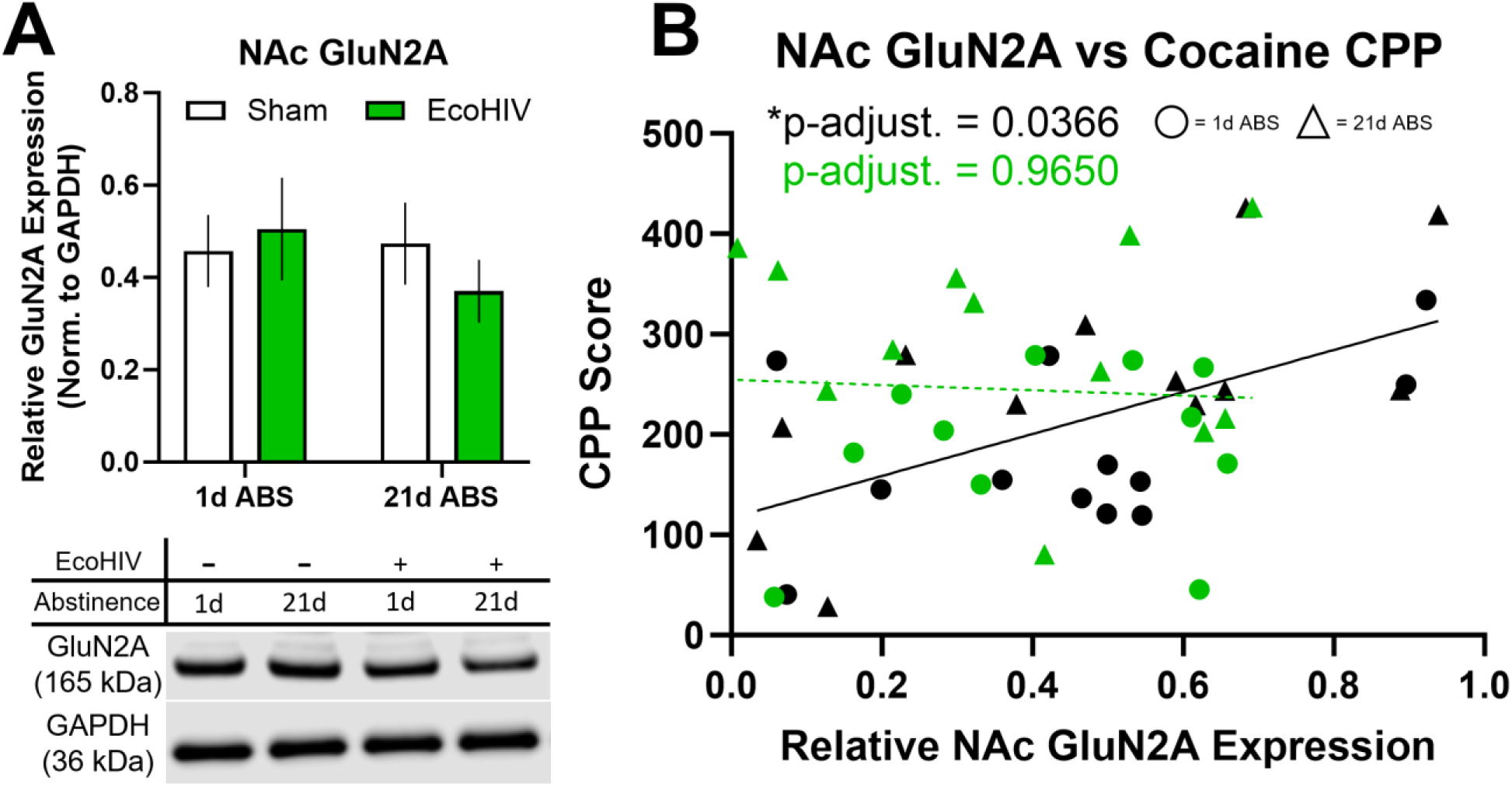
NAc GluN2A expression and associated cocaine seeking following daily conditioning and abstinence. (A) Among mice conditioned daily, no group mean differences in NAc GluN2A expression were detected due to EcoHIV, abstinence, or their interaction. Error bars = ±SEM; n = 11-12/group. (**B**) Cocaine seeking behavior was significantly correlated with GluN2A expression only among sham mice across abstinence time points.

### Peripheral cytokine, chemokine, and growth factor expression

In addition to exploring corticolimbic neuroimmune and glutamatergic mechanisms that may underlie EcoHIV-induced impairments in cocaine seeking, we sought to characterize changes in peripheral immune function due to EcoHIV infection, cocaine abstinence, and cocaine exposure pattern. Here, we used a multiplex array on plasma samples collected at final tissue collection to assess cytokine, chemokine, and growth factor expression in an exploratory manner. Immune marker levels were analyzed via one-way ANOVAs for each experiment, and analyses that survived p-value correction for multiple testing were further probed for *post hoc* comparisons of each experimental group to a reference control group of sham mice that received only saline injections via Dunnett’s multiple comparison testing. Among mice in Experiment 1, a Welch’s one-way ANOVA revealed significant group differences in granulocyte-colony stimulating factor (G-CSF) expression (**Figure 6A**; *F*_(4,23.59)_ = 8.046, p = 0.0003, p-adj. = 0.0045; 2 outliers in the EcoHIV-21d ABS group were excluded). *Post hoc* analysis revealed that at 1d ABS, both sham- and EcoHIV-infected mice exhibited decreased G-CSF expression relative to the reference control group (*p < 0.01). In addition to G-CSF, a Welch’s one-way ANOVA reveled significant group differences in vascular endothelial growth factor (VEGF) expression (**Figure 6B**; *F*_(4,24.58)_ = 4.593, p = 0.0032, p-adj. = 0.0439; 1 outlier in the EcoHIV-1d ABS group and 1 outlier in the Sham-1d ABS group were excluded). *Post hoc* analysis did not reveal significant group differences, although a trend for a decrease in EcoHIV-1d ABS mice relative to the reference control group was noted (p = 0.0848). Among mice in Experiment 2, a Welch’s one-way ANOVA revealed significant group differences in interleukin (IL)-5 expression (**Figure 7**; *F*_(4,25.35)_ = 6.542, p = 0.0009, p-adj. = 0.0134; 1 outlier in the Control group and 1 outlier in the EcoHIV-1d ABS group were excluded). *Post hoc* analysis revealed that a significant decrease in IL-5 in Sham-1d ABS mice (*p < 0.05) and a trend towards decreased IL-5 in EcoHIV-1d ABS mice (p < 0.10) relative to the reference control group. Altogether, these results indicate that the expression of specific peripheral immune markers was generally suppressed at 1d ABS following cocaine exposure, and that the profile of such deficits depends on the cocaine exposure pattern.

**Figure 6.**
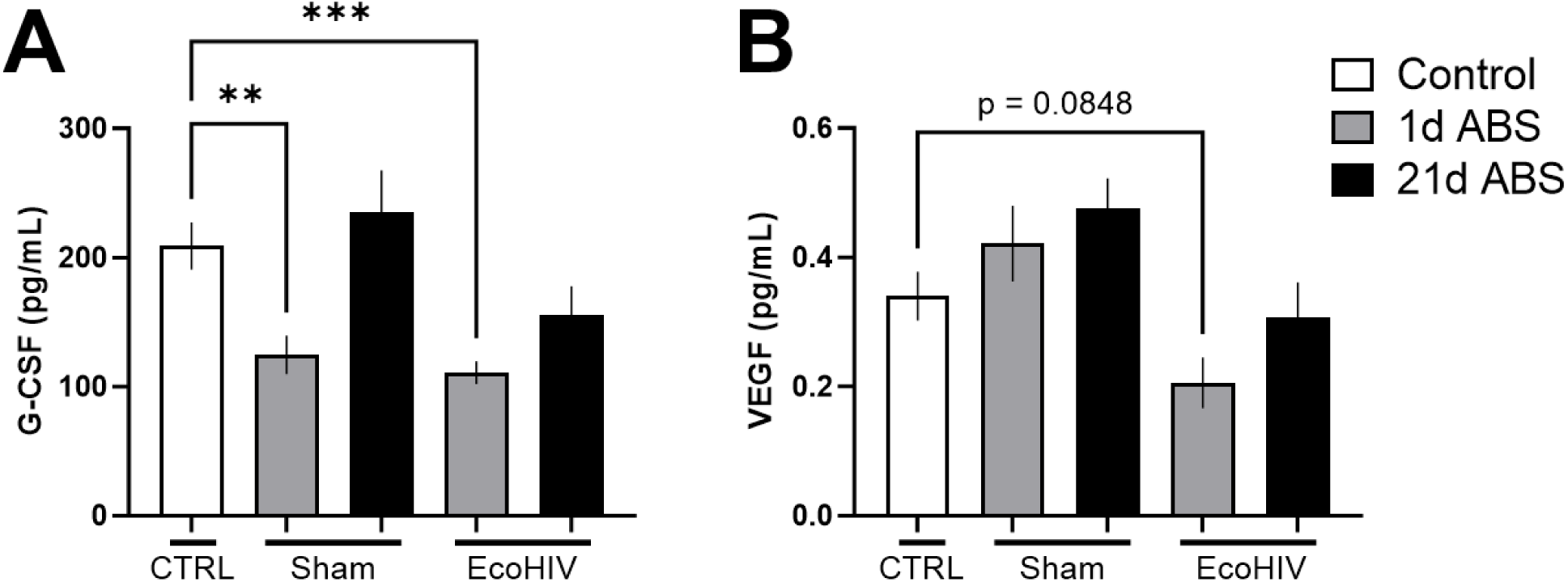
Plasma G-CSF and VEGF expression following daily cocaine exposure. Multiplex cytokine arrays were used to explore peripheral cytokine, chemokine, and growth factor expression in plasma following daily cocaine conditioning and abstinence. Plasma immune factor levels across treatment groups were compared to a reference control group of sham-infected mice that received only saline injections (i.e., “CTRL”). (**A**) Following 1d ABS, both sham- and EcoHIV-infected mice exhibited significantly reduced granulocyte-colony stimulating factor (G-CSF) expression relative to the reference control group (**p < 0.01;***p < 0.001; n = 10-12/group). (**B**) EcoHIV-infected mice showed a trend towards reduced vascular endothelial growth factor (VEGF) expression relative to the reference control group (n = 10-12/group). Error bars = ±SEM.

**Figure 7.**
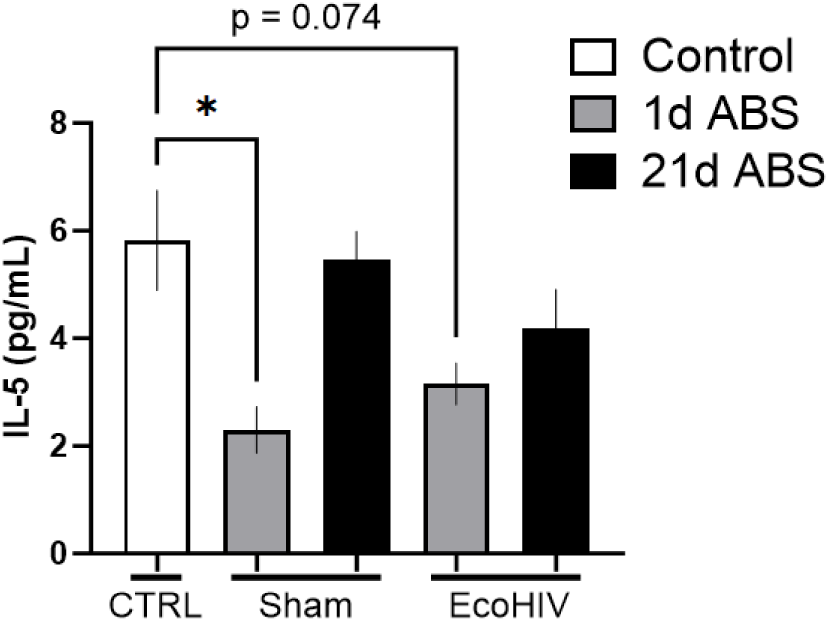
Plasma IL-5 expression following intermittent cocaine exposure. Multiplex cytokine arrays were used to explore peripheral cytokine, chemokine, and growth factor expression in plasma following intermittent cocaine conditioning and abstinence. Plasma immune factor levels across treatment groups were compared to a reference control group of sham-infected mice that received only saline injections (i.e., “CTRL”). 1d ABS resulted in a significant reduction (*p < 0.05), and a trend towards a reduction, in IL-5 expression for sham- and EcoHIV-infected mice, respectively. n = 10-12/group. Error bars = ±SEM.

## Discussion

Comorbidities that often accompany substance use disorders (SUDs), including HIV, can substantially alter the biobehavioral landscape that characterizes each disease state (Namba et al., 2021). While many studies have demonstrated the deleterious effects of chronic drug exposure on the pathophysiology of HIV, relatively little is known about how chronic HIV infection alters addiction-related outcomes. This represents a critical gap in the field, given that successful treatment of SUDs for PLWH could also provide substantial improvements to the long-term clinical management of HIV (Durvasula and Miller, 2014; Dash et al., 2015; Kumar et al., 2015; Bertholet et al., 2023). Thus, in the present study, we sought to determine the interactive effects of EcoHIV infection, cocaine exposure pattern, and cocaine abstinence on subsequent cocaine-seeking behavior and associated neuroimmune outcomes.

Protracted abstinence from drug use represents a significant relapse risk factor, where drug-associated cues and environmental stimuli can trigger craving and subsequent relapse (Gawin and Kleber, 1986; Carter and Tiffany, 1999; Parvaz et al., 2016). This abstinence-dependent process, termed incubation of craving (Grimm et al., 2001), occurs across drug classes (Pickens et al., 2011; Li et al., 2015). However, until the present study, understanding of the impact of chronic HIV infection on this process across different cocaine exposure patterns was limited. One key finding presented here is that the impact of EcoHIV infection on the incubation effect depended on the cocaine exposure pattern. This was paralleled by distinct neurobiological and peripheral immune adaptations, suggesting that treatment targets identified for SUD medications development should critically consider factors such as drug use patterns and comorbidity status. Taken together, our findings underscore the need for more holistic perspectives when modeling SUDs preclinically, especially as it pertains to medications development.

### The impact of EcoHIV infection on incubated cocaine seeking depends on cocaine exposure pattern

We found that the effect of EcoHIV infection on the incubation of cocaine seeking following a period of protracted abstinence in male mice depended on the cocaine exposure pattern during cocaine conditioning. Sham-infected mice receiving four daily cocaine conditioning sessions exhibited significant incubation of cocaine seeking across the CPP test session after 21 days of forced abstinence, while sham-infected mice receiving four cocaine conditioning sessions in between saline conditioning sessions (i.e., intermittent conditioning) exhibited a strong trend towards incubation of cocaine CPP. Among EcoHIV-infected mice, daily conditioning produced significant incubation after both 1 and 21 days of abstinence, while intermittent conditioning produced the reverse pattern of abstinent-dependent cocaine seeking, with higher cocaine CPP at 1 day abstinence. These differences in behavior between these two drug exposure regimens were evident in the cocaine-induced locomotion data collected during cocaine conditioning, where the interaction between EcoHIV infection and cocaine on locomotor behavior differed substantially between these two paradigms (Figures 2A,C). Specifically, EcoHIV-infected mice showed a more robust sensitization to the locomotion-enhancing effects of cocaine when they were conditioned daily as opposed to intermittently. Despite behavioral differences, we did not identify any significant changes in glutamate receptor expression or fractalkine expression within the mPFC and NAc in mice exposed intermittently to cocaine. However, given the locomotor behavior data, it is possible that the effects of combined EcoHIV infection and varying cocaine exposure patterns could produce distinct adaptations in other systems, such as in dopamine transmission, that could underlie the observed differences in cocaine-induced locomotion.

Within the cocaine self-administration literature, studies suggest that intermittent access to intravenous cocaine, as opposed to continuous access, results in greater cocaine-induced dopamine release and reinforcing efficacy of cocaine, particularly after a short abstinence period (Calipari et al., 2014; Kawa et al., 2019b, 2019a). Such studies make clear that the temporal pattern of drug exposure is a critical variable that determines drug-induced dopaminergic plasticity. Complementing this is the known temporal dynamics of drug abstinence effects on subsequent relapse-like behavior, where the peak levels of drug seeking (or craving as reported in humans) varies depending on the drug and, evidently, the temporal pattern of drug exposure. For example, one recent study showed an overall greater incubation effect following intermittent access compared to a continuous access schedule of intravenous cocaine self-administration, but only 2 and 29 day abstinence timepoints were examined (Nicolas et al., 2019). As we only examined abstinence days 1 and 21 in the present study, an intriguing possibility that warrants further investigation is whether intermittent cocaine exposure, as described in this study, might result in a shift in peak cocaine seeking among EcoHIV-infected mice to an earlier time point (e.g., 7 days). Interestingly, one longitudinal study reported that PLWH who inject drugs exhibited a shorter time to first relapse following initial cessation of injection drug use (Shah et al., 2006), which highlights the possibility of alternative incubation of craving processes among PLWH. Thus, future studies should probe alternative relapse timepoints to more fully characterize changes in peak cocaine seeking between different cocaine exposure conditions.

The present results replicate findings from another CPP study demonstrating the incubation of cocaine seeking (Lubbers et al., 2016). Indeed, this would be concordant with the extant literature on the incubation of cocaine seeking following intravenous cocaine self-administration (Tran-Nguyen et al., 1998; Grimm et al., 2001, 2003; Thiel et al., 2012; Ma et al., 2014). However, a novel finding here that adds to this existing literature is that both intermittent and daily cocaine conditioning recapitulate this incubation effect in sham control animals, with a more robust incubation effect following daily conditioning. Unexpectedly, the effect of EcoHIV on this critical addiction-related behavioral process was contingent on the cocaine conditioning regimen. Extrapolating out to the clinical population, such findings would suggest that consideration of drug use patterns may be warranted to achieve successful SUD treatment outcomes for PLWH. This is particularly important considering that PLWH tend to report higher rates of drug use compared to the general population (Shiau et al., 2017). Nevertheless, it is important to note that only male mice were tested here, and the results may therefore be restricted to only males. Indeed, there are sex differences in the temporal pattern of cocaine seeking across abstinence, as well as associated corticolimbic cellular adaptations (Kim et al., 2022; Towers et al., 2023), though this may depend on estrous phase (Corbett et al., 2021). Thus, examination of sex differences in cocaine seeking across the various factors and levels of analysis tested here represents an important future direction of this work.

### Spleen expression of HIV-1 DNA depends on cocaine exposure pattern

To validate the infection status of mice in this study, we measured levels of HIV-1 LTR DNA from spleen tissue isolates (Figure 3). We then compared these DNA levels across abstinence and cocaine exposure pattern conditions, where we observed a significant increase in HIV-1 LTR DNA levels in mice following daily conditioning and 1d ABS. By 21d ABS in these mice, LTR DNA levels were equal to those observed at both abstinence timepoints in mice exposed to cocaine intermittently. Interestingly, cocaine is known to promote HIV transcriptional initiation through activation of NF-κB and subsequent interactions with the LTR (Sahu et al., 2015; Tyagi et al., 2016). Thus, higher levels of LTR DNA following repeated, continuous cocaine exposure highlights the potential for increased vulnerability to transcription of viral substrates. The LTR (particularly the 5’ LTR) is a critical coding region of the viral genome that is necessary for transcription of viral RNA and, ultimately, the synthesis of new viral particles (Klaver and Berkhout, 1994; Krebs et al., 2001). It is important to note that the qPCR methodology used in this study cannot differentiate between integrated vs. non-integrated DNA. During the asymptomatic phase of HIV infection, full-length, linear, unintegrated DNA represents the vast majority of total viral DNA (Chun et al., 1997), and these high levels of unintegrated viral DNA can be found in both brain and lymphoid tissues (Pang et al., 1990; Chun et al., 1997; Teo et al., 1997). Interestingly, several studies suggest that at least some viral proteins, such as Nef and Tat, can be synthesized from unintegrated viral DNA (for review, see Wu, 2004). In EcoHIV-infected mice, high viremia is observed 1-week post-inoculation, which drops precipitously to near-undetectable levels by weeks 3-5 post-inoculation (Gu et al., 2018; Alfar et al., 2023). However, both total and integrated viral DNA levels remain persistently elevated and stable for weeks to months post-inoculation in the spleen (Gu et al., 2018). Taken altogether, it is possible that unintegrated viral DNA that persists in lymphoid (and possibly brain) tissue may contribute to chronic, low-level exposure to viral proteins that could impact nervous system function in the absence of active transcription of integrated proviral DNA and subsequent production of viral particles. This highlights the significance of our finding of increased splenic HIV-1 LTR DNA following daily conditioning and acute abstinence and points towards a potential mechanism through which cocaine and subsequent abstinence impact the pathophysiology of HIV and, ultimately, neurobehavioral impairment.

### mPFC fractalkine expression is associated with cocaine-seeking behavior

As the primary CNS reservoir for HIV, microglial dysregulation is putatively implicated in contributing to HIV-associated neurocognitive impairments. Indeed, an extensive body of literature indicates microglia as key regulators of synaptic plasticity (Ben Achour and Pascual, 2010; Kovács, 2012; Levin and Godukhin, 2017; Cornell et al., 2022), and one key molecular mediator of neuron-microglia interactions is fractalkine signaling (Pawelec et al., 2020). Thus, we examined CX3CL1 (i.e., fractalkine) and CX3CR1 expression within the mPFC and NAc and assessed whether the expression of these substrates correlated with cocaine seeking. Within the mPFC of mice conditioned daily, mean fractalkine expression was not different across treatment groups (Figure 4A), but significantly correlated with cocaine CPP among sham mice (Figure 4B). Moreover, no such effects were observed for mice conditioned intermittently, which corroborates a recent report showing no effect of intermittent cocaine conditioning on PFC fractalkine expression (Rosa et al., 2022). Fractalkine signaling can regulate microglial responses to inflammatory insults, including exposure to drugs and HIV-1 proteins (Cotter et al., 2002; Namba et al., 2021). Specifically, fractalkine signaling can serve as an “off signal” for microglial activation (Pawelec et al., 2020), which could protect neuronal circuits and synapses from microglia activation-induced immunomodulation. However, as is the case for many cytokines and chemokines, their role in regulating synaptic plasticity and subsequent behavior is likely brain region- and disease state-specific (Li et al., 1997; Schneider et al., 1998; Jankowsky et al., 2000; Beattie et al., 2002; Stellwagen et al., 2005; Yang et al., 2005; Lewitus et al., 2014, 2016; Pawelec et al., 2020).

Very few *in vivo* studies have demonstrated HIV and cocaine abstinence effects on fractalkine signaling within the mPFC, although *in vitro* studies may provide useful insights into the role of *in vivo* fractalkine signaling in regulating drug seeking. For example, co-exposure of CX3CR1-deficient striatal neurons *in vitro* to Tat and morphine synergistically enhances microglial motility, dendritic pruning, and cell death, and exogenous fractalkine is protective against such effects in wild type neurons (Suzuki et al., 2011). Microglia exposed *in vitro* to HIV-1 Tat also exhibit impaired CX3CR1 expression, which is driven by nuclear factor kappa B (NF-κB) pathway signaling (Duan et al., 2014). Importantly, we and others have shown that striatal NF-κB mediates relapse-like drug seeking in rodents (Russo et al., 2009; Namba et al., 2020, 2022), highlighting the *in vivo* translatability of these *in vitro* studies. We have also recently shown that NAc core (NAcore) fractalkine expression is negatively correlated with cue-induced cocaine seeking in rats (Namba et al., 2023a). Thus, if fractalkine is serving a similar protective role within the mPFC, the positive correlation between mPFC fractalkine expression and cocaine seeking in sham mice could represent a behavior-dependent compensatory mechanism that helps suppress cocaine seeking.

A crucial behavioral consideration here is that mice in this study were tested for CPP in a drug-free state, which is functionally an extinction learning setting. A recent study showed that CX3CR1-KO mice show significant extinction learning impairments in a cocaine CPP paradigm, while acquisition and reinstatement remained intact, and an acute injection of cocaine in wild-type mice with a history of repeated exposure to cocaine resulted in enhanced PFC expression of fractalkine (Montesinos et al., 2020). Interestingly, we recently have shown that HIV-1-infected humanized mice exhibit impaired extinction of cocaine seeking (Buck et al., 2024), a PFC-dependent behavior (Peters et al., 2008; Lalumiere et al., 2012; Nett et al., 2023). Taken altogether, the positive relationship between mPFC fractalkine expression and cocaine seeking in sham- but not EcoHIV-infected mice observed in our work (Figure 4B) is consistent with a compensatory feedback mechanism to regulate extinction learning, which may occur in an activity-dependent manner that involves transient release of fractalkine expression and may be impaired in EcoHIV-infected mice.

### NAc GluN2A expression is associated with cocaine-seeking behavior

Akin to the observed effect with fractalkine, a significant positive relationship was observed between NAc GluN2A expression and cocaine CPP in sham mice that received daily cocaine conditioning (Figure 5B). While the lack of an overall mean difference in total GluN2A protein expression within the NAc among sham animals (Figure 5A) is congruent with a previous CPP study in rats (Huang et al., 2009), the significant correlation between GluN2A expression and CPP reported here suggests that individual variability in behavior may be predicted by NAc GluN2A expression despite no overall population differences across treatment groups. It is important to note that the tissue isolates in this study were prepared as whole cell lysates for western blot analysis. Crude membrane fractions or synaptosome preparations may have revealed more nuanced and functionally relevant differences in the expression levels of both NMDA and AMPA receptor subunits. For example, a recent study showed that GluN2A expression is increased only in postsynaptic density fractions of the prelimbic cortex and dorsal hippocampus of rats with a history of cocaine self-administration and extinction (Smaga et al., 2021). Regardless, the present findings suggest that NAc GluN2A expression depends on the temporal dynamics of cocaine exposure and HIV infection status.

GluN2A-containing NMDARs exhibit faster decay kinetics and lower binding affinity for downstream calcium/calmodulin kinase II (CAMKII) than GluN2B-containing NMDARs (Cull-Candy and Leszkiewicz, 2004). Thus, it is hypothesized that increased synaptic insertion of GluN2B NMDARs, rather than GluN2A NMDARs, confers greater potential for sustained postsynaptic excitation of medium spiny neurons (MSNs) within the NAc and subsequent drug-seeking behavior. Indeed, several studies have established a role for NAc expression of GluN2B in mediating drug relapse-like behavior (Huang et al., 2009; Lee and Dong, 2011; Shen et al., 2011; Gipson et al., 2013). These findings led us to hypothesize that EcoHIV-induced increases in cocaine seeking would be associated with enhanced NAc GluN2B expression. Unexpectedly, we did not observe any significant effect across treatment groups for GluN2B expression and observed a significant correlation between cocaine seeking and GluN2A expression. Nevertheless, increased NAc expression of GluN2A may still confer greater risk for drug seeking. For example, protracted withdrawal with extinction training following nicotine self-administration in rats is associated with increased postsynaptic excitability within NAc MSNs, which is accompanied by increased expression of both GluN2A and GluN2B (in a crude membrane fraction) and transient, nicotine seeking-dependent increases in dendritic spine head diameter (Gipson et al., 2013). Importantly, *in vivo* pharmacological inhibition of GluN2A via intra-NAc TCN-201 treatment dose-dependently suppressed nicotine seeking in this study. Intra-NAc treatment with the GluN2A antagonist NVP can also decrease cocaine-seeking behavior under extinction conditions (Hafenbreidel et al., 2017). Altogether, these studies indicate that increased NAc GluN2A expression may confer increased vulnerability to relapse-like drug seeking.

The positive correlation between NAc GluN2A expression and cocaine seeking among sham mice in this study, and only among those conditioned daily, is a unique finding that converges onto existing literature that implicates GluN2A in drug relapse-like behavior and associated synaptic plasticity. Moreover, the lack of a correlation between cocaine seeking in EcoHIV-infected mice and GluN2A (among the other substrates measured) highlights the need to consider alternative corticostriatal mechanisms that may facilitate the potentiated cocaine seeking we observed in these mice, including HIV-induced dysregulation of corticostriatal synaptic architecture (McLaurin et al., 2018, 2019, 2022).

### Cocaine exposure pattern impacts the effect of EcoHIV on peripheral immune function

Cocaine exposure and withdrawal disrupt both central and peripheral immune function. For example, plasma cortisone levels and peripheral lymphocyte proliferation responses in rats are suppressed for up to 6 days of withdrawal (but not at later withdrawal timepoints) following repeated exposure to 10 mg/kg cocaine (Avila et al., 2003). Our findings have shown in rats that 5 days of abstinence from cocaine self-administration and/or exposure to the HIV protein gp120 is associated with the suppression of numerous cytokines and chemokines within the NAc, including fractalkine, IL-4, IL-5, IL-18, and several others (Namba et al., 2023a). This work expands on previous findings by elucidating whether the impact of cocaine abstinence on immune outcomes, and the potential modulatory effect of EcoHIV infection, would depend on the cocaine exposure pattern. Here, we demonstrate differential effects of cocaine abstinence and EcoHIV infection on plasma immune markers between intermittent and daily cocaine exposure. Specifically, 1d ABS from intermittent cocaine conditioning produced a significant reduction in plasma IL-5 expression in sham mice, with a similar trend observed for EcoHIV-infected mice (Figure 6). A different profile of peripheral immune response was observed among mice conditioned daily with cocaine. Specifically, G-CSF was significantly suppressed at 1d ABS for both sham- and EcoHIV-infected mice (Figure 7A), with a trend towards suppressed VEGF expression in EcoHIV-infected mice at 1d ABS.

Many studies have demonstrated that the balance of T helper 1 (Th1) and Th2 cytokine profiles are dysregulated after chronic cocaine exposure and subsequent withdrawal (Stanulis et al., 1997; Levandowski et al., 2016; Zaparte et al., 2019). Moreover, the levels of Th1, Th2, and Th17 cytokines change over the course of HIV infection, from acute infection to viral suppression via ART, and the interactions between these different immune substrates ultimately dictates the fate of cellular-mediated immunity and disease progression (Kedzierska and Crowe, 2001). A recent study examining plasma cytokine, chemokine, and growth factor changes during primary HIV infection (prior to ART therapy) demonstrated increases in several targets in plasma, notably including IL-5 and G-CSF. In particular, G-CSF was negatively correlated with CD4/CD8 T cell ratio (Bordoni et al., 2020). In the EcoHIV model, mice exhibit an inherent antiviral immune response that leads to viral suppression in the absence of ART (Gu et al., 2018). While this may represent a useful model of virally-suppressed PLWH, it remains a limitation to be noted. Thus, examination of cytokine expression at an earlier timepoint post-inoculation may have recapitulated the immune profiles observed among clinical populations during this critical window of HIV-1 disease progression.

Our findings here are in accordance with studies that implicate early abstinence and withdrawal from cocaine as a sensitive period of immune vulnerability. Notably, we did not observe any significant differences in cytokine, chemokine, or growth factor expression following 21d ABS from cocaine compared to the sham-saline control group, indicating that cocaine-induced impairments in peripheral immune signaling are likely transient in a subchronic exposure model. However, early withdrawal from cocaine may represent a period of increased vulnerability to infection by pathogens like HIV (Avila et al., 2003, 2004). We further did not observe any significant effects of EcoHIV infection on any of these measures. This is likely due to the fact that EcoHIV-infected animals are not viremic and are otherwise virally suppressed by the time cocaine conditioning (and subsequent CPP testing and tissue collection) occurs (Gu et al., 2018).

## Conclusion

Current findings suggest that the impact of EcoHIV infection on cocaine-seeking behavior is accompanied by central nervous system impairments, as opposed to solely peripheral immune dysfunction, in a manner that is sensitive to the temporal dynamics of drug exposure and subsequent abstinence. We hypothesize that corticolimbic circuit mechanisms not explored in this study may mediate the EcoHIV-induced deficits in cocaine seeking. As demonstrated in other HIV rodent models, chronic exposure to viral proteins can disrupt corticostriatal dendritic spine morphology (McLaurin et al., 2018, 2019, 2022). This could perhaps alter the ratio of dopaminergic and glutamatergic synaptic input onto MSNs within the NAc, which make symmetric and asymmetric contact with dendritic spines, respectively (Freund et al., 1984; Spiga et al., 2014). Such disruptions in synaptic architecture could drive HIV-induced dysregulations in reward seeking. While unexplored in this study, these alternative mechanisms warrant future investigation, particularly in the context of incubated cocaine craving and across various drug exposure regimens.

Taken altogether, this study underscores the importance of factors such as drug use patterns and abstinence as key factors that may determine not only the influence of HIV on addiction-related outcomes, but also have implications for the pathophysiology of HIV itself. Thus, our findings support the broader position that proper long-term clinical management of HIV among people who use drugs must consider the treatment of SUDs in a more holistic manner, taking into consideration factors such as abstinence and temporal patterns of drug use.

## Acknowledgements

We would like to thank Christine Side and Lauren Buck of the Barker Lab for their excellent technical expertise. We also thank the Volsky Laboratory (Icahn School of Medicine at Mount Sinai) for generously providing us with EcoHIV constructs. These studies were supported by NIH grants DP2DA051907 and DP2DA051907-01S1 (JMB) and a pilot grant awarded to MDN from the Drexel University-Temple University Comprehensive NeuroHIV Center (NIH grant P30MH0921777). Figure 1 was prepared using Biorender.com.

